# Regulation by Progestins, Corticosteroids and RU486 of Activation of Elephant Shark and Human Progesterone Receptors: An Evolutionary Perspective

**DOI:** 10.1101/2021.01.20.427507

**Authors:** Xiaozhi Lin, Wataru Takagi, Susumu Hyodo, Shigeho Ijiri, Yoshinao Katsu, Michael E. Baker

**Affiliations:** Graduate School of Life Science, Hokkaido University, Sapporo, Japan; Laboratory of Physiology, Atmosphere and Ocean Research Institute, University of Tokyo, Chiba, Japan; Graduate School of Fisheries Science, Hokkaido University, Hakodate, Japan; Faculty of Science, Hokkaido University, Sapporo, Japan; Division of Nephrology, Department of Medicine, University of California, San Diego, CA, USA; Center for Academic Research and Training in Anthropogeny (CARTA), University of California, San Diego, CA, USA

**Keywords:** elephant shark PR, progesterone receptor evolution, progestins, corticosteroids, RU486

## Abstract

We investigated progestin and corticosteroid activation of the progesterone receptor (PR) from elephant shark, a cartilaginous fish belonging to the oldest group of jawed vertebrates. Comparison with human PR provides insights into the evolution of steroid activation of human PR. At 1 nM steroid, elephant shark PR is activated by progesterone, 17-hydroxy-progesterone, 20β-hydroxy-progesterone, 11-deoxycorticosterone (21-hydroxyprogesterone) and 11-deoxycortisol. Human PR, in comparison, is activated at 1 nM steroid, only by progesterone and 11-deoxycorticosterone, indicating increased progestin and corticosteroid specificity during the evolution of human PR. RU486, an important clinical antagonist of human PR, did not inhibit progesterone activation of elephant shark PR. Cys-528 in elephant shark PR corresponds to Gly-722 in human PR, which is essential for RU486 inhibition of human PR. Confirming the importance of Cys-528 in elephant shark PR, RU486 inhibited progesterone activation of the Cys528Gly mutant PR. Compared to wild-type human PR, there was an increase in activation of human Gly722Cys PR by11-deoxycortisol and a decrease in activation by corticosterone, which may have been important in selection for the mutation corresponding to human glycine-722 PR that first evolved in platypus PR, a basal mammal.

## Introduction

The progesterone receptor (PR) receptor belongs to the nuclear receptor family, a diverse group of transcription factors that also includes the glucocorticoid receptor (GR), mineralocorticoid receptor, androgen receptor (AR), and estrogen receptor (ER) (1–3). In humans, the progesterone receptor (PR) mediates progesterone regulation of female reproductive physiology in the uterus and mammary gland, including fertilization, maintenance of pregnancy and preparation of the endometrium for implantation and parturition (4–7). Moreover, progesterone has important physiological actions in males, including in the prostate and testes (8–11). Further, progesterone activates the PR in the brain, bone, thymus, lung and vasculature in females and males (12,13). Thus, progesterone is a steroid with diverse physiological activities in many organs in females and males.

Although activation by progesterone of the PR in chickens (4,14), humans (15), and zebrafish (16,17) has been examined, steroid activation of a PR in the more basal cartilaginous fish lineage has not been fully investigated. To remedy this omission, we studied the activation by a panel of progestins and corticosteroids (Figure 1) of the PR from the elephant shark (*Callorhinchus milii*), a cartilaginous fish belonging to the oldest group of jawed vertebrates, which diverged about 450 million years ago from bony vertebrates (18,19).

**Figure 1.**
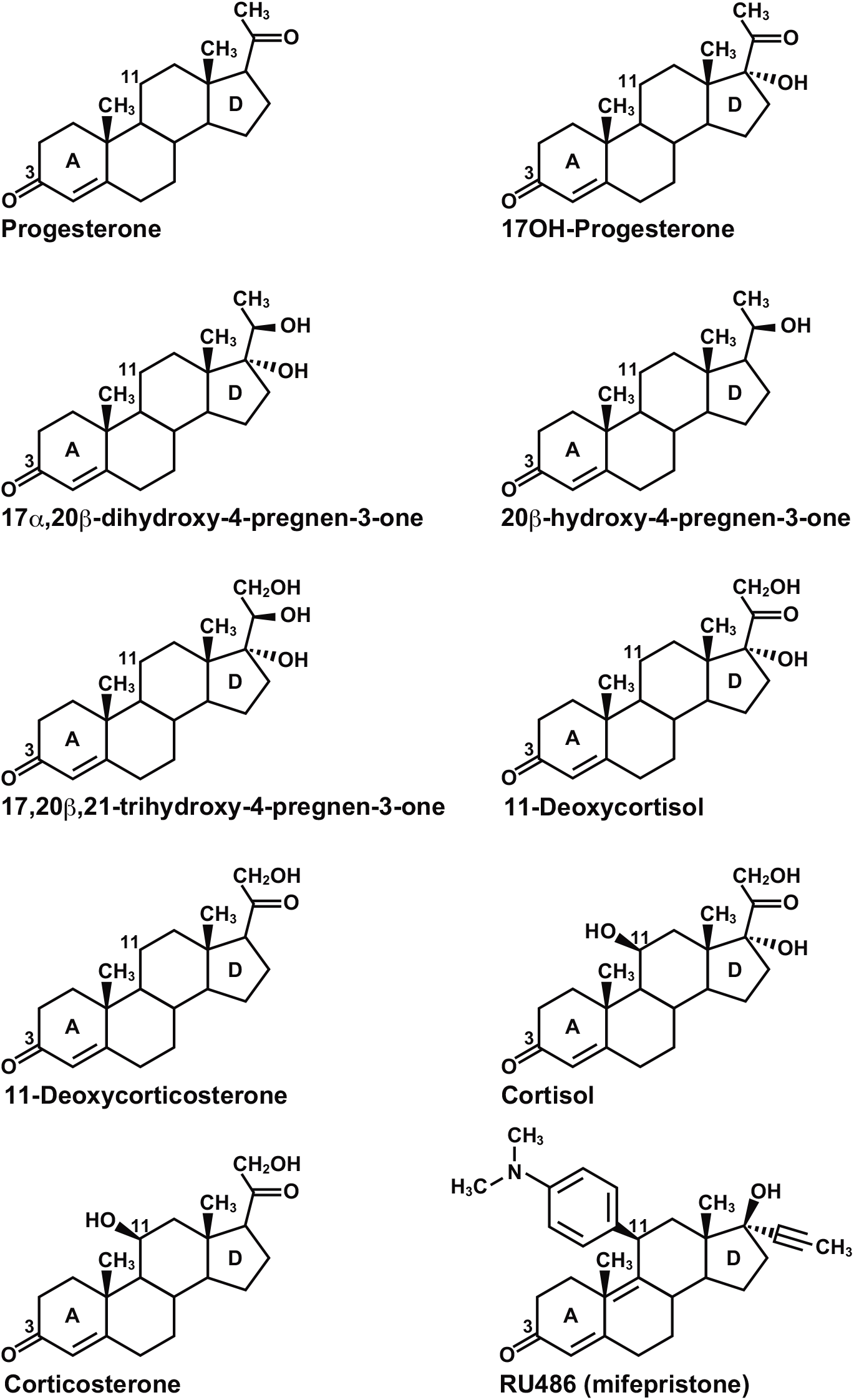
Structures of corticosteroids and progestins. Progesterone is female reproductive steroid that also is important in male physiology (4,11). 17,20β-dihydroxy-progesterone is a maturation inducing hormone of teleost fish (20–22). 17,20β,21-trihydroxy-progesterone is a major ovarian steroid produced by the teleost fish (23). Cortisol, corticosterone and 11-deoxycortisol are physiological glucocorticoids in terrestrial vertebrates and ray-finned fish (24,25). 11-deoxycorticosterone is a mineralocorticoid (25–28). RU486 is an antagonist of human PR (29,30).

Elephant shark PR is an attractive receptor to investigate the ancestral regulation of steroid-mediated PR transcription because, in addition to its phylogenetic position as a member of the oldest lineage of jawed vertebrates, genomic analyses reveal that elephant shark genes are evolving slowly (19), making studies of its PR useful for studying ancestral proteins, including the PR, for comparison for similarities and differences with human PR to elucidate the evolution of steroid specificity for the PR in terrestrial vertebrates (19,31,32). In addition, we were interested in the response of elephant shark PR to RU486 (Mifepristone), which is an antagonist for the human PR (29,30,33) and also a potential anticancer drug for treating progesterone-dependent breast cancer (34).

We find that elephant shark PR is activated by progesterone, 17-hydroxy-progesterone, 20β-hydroxy-progesterone, 17,20β-dihydroxy-progesterone, corticosterone, 11-deoxycorticosterone (21-hydroxy-progesterone) and 11-deoxycortisol. In contrast human PR is activated only by progesterone, 20β-hydroxy-progesterone, 11-deoxycorticosterone and corticosterone, indicating that human PR has increased specificity for progestins and corticosteroids.

We also find that RU486 does not inhibit progesterone activation of elephant shark PR. We show that this is due to cysteine-528 in elephant shark PR, which corresponds to glycine-722 on human PR, an amino acid that Benhamou et al. (35) reported was essential for antagonist activity of RU486. They found that mutation of glycine-722 to cysteine abolished RU486 inhibition of progesterone activation of human PR. Analyses of vertebrate PRs reveals that an ancestor of human PR-Gly722 first appeared in platypus, a basal mammal (31).

To search for functional changes in human PR that correlate with the evolution of RU486 antagonist activity, we constructed the elephant shark PR-Gly528 mutant and the human PR-Cys722 mutant and studied their activation by several steroids. Elephant shark PR-Gly528 had a weaker response to 11-deoxycortisol and 17-hydroxy-progesterone. Human PR-Cys722 displayed increased activation by 11-deoxycortisol and decreased activation by corticosterone. An altered response to one or more of these steroids may have been selective for the evolution of an ancestor of glycine-722 in a PR in an ancestral platypus at the base of the mammalian line.

Lastly, our studies provide an insight into the evolution of steroid activation of fish PRs. In fish, instead of progesterone, it is 17,20β-dihydroxy-progesterone that is the physiological ligand for the PR in zebrafish (16,17) and other teleosts (20–22,36,37). We find that for elephant shark PR, the half-maximal response (EC50) of progesterone and 17,20β-dihydroxy-progesterone are 0.18 nM and 2.6 nM, respectively. This ten-fold higher EC50 of elephant shark PR for 17,20β-dihydroxy-progesterone compared to progesterone, indicates that the during the evolution of ray-finned fish, there was a reversal between progesterone and 17,20β-dihydroxy-progesterone in their selectivity for elephant shark PR and for zebrafish PR and other fish PRs, and that this role for 17,20β-dihydroxy-progesterone, instead of progesterone, as a ligand for fish PR evolved after the divergence of ray-finned fish from cartilaginous fish (18,19).

## Materials and Methods

### Chemical reagents

Cortisol, corticosterone, 11-deoxycorticosterone, 11-deoxycortisol, progesterone, 17α-hydroxy-progesterone, 17,20β,21-tri-hydroxy-progesterone, 20β-hydroxy-progesterone, and 17,20β-dihydroxy-progesterone were purchased from Sigma-Aldrich. RU486 was purchased from Cayman Chemical. For reporter gene assays, all hormones were dissolved in dimethyl-sulfoxide (DMSO); the final DMSO concentration in the culture medium did not exceed 0.1%.

### Construction of plasmid vectors

The full-length PRs were amplified by PCR with KOD DNA polymerase. The PCR products were gel-purified and ligated into pcDNA3.1 vector (Invitrogen). Site-directed mutagenesis was performed using KOD-Plus-mutagenesis kit (TOYOBO). All cloned DNA sequences were verified by sequencing.

### Transactivation assay and statistical methods

Transfection and reporter assays were carried out in HEK293 cells, as described previously (38,39). All experiments were performed in triplicate. The values shown are mean ± SEM from three separate experiments, and dose-response data, which were used to calculate the half maximal response (EC50) for each steroid, were analyzed using GraphPad Prism. Comparisons between two groups were performed using paired *t*-test. *P* < 0.05 was considered statistically significant. The use of HEK293 cells and an assay temperature of 37C does not replicate the physiological environment of elephant sharks. Nevertheless, studies with HEK293 cells and other mammalian cell lines have proven useful for other studies of transcriptional activation by steroids of steroid hormone receptors from non-mammalian species (39–41).

## Results

### Transcriptional activation of full-length elephant shark PR by progestins and corticosteroids

We screened a panel of steroids for transcriptional activation of full-length elephant shark and human PRs using HEK293 cells. At 10 nM, progesterone, 17-hydroxy-progesterone, 17, 20β-dihydroxy-progesterone, a fish maturation hormone, and 20β-hydroxy-progesterone activated elephant shark PR (Figure 3A). At 10 nM, 11-deoxycorticosterone (21-hydroxyprogesterone), 11-deoxycortisol and corticosterone activated elephant shark PR (Figure 3C) indicating that elephant shark PR responds to corticosteroids.

**Figure 3.**
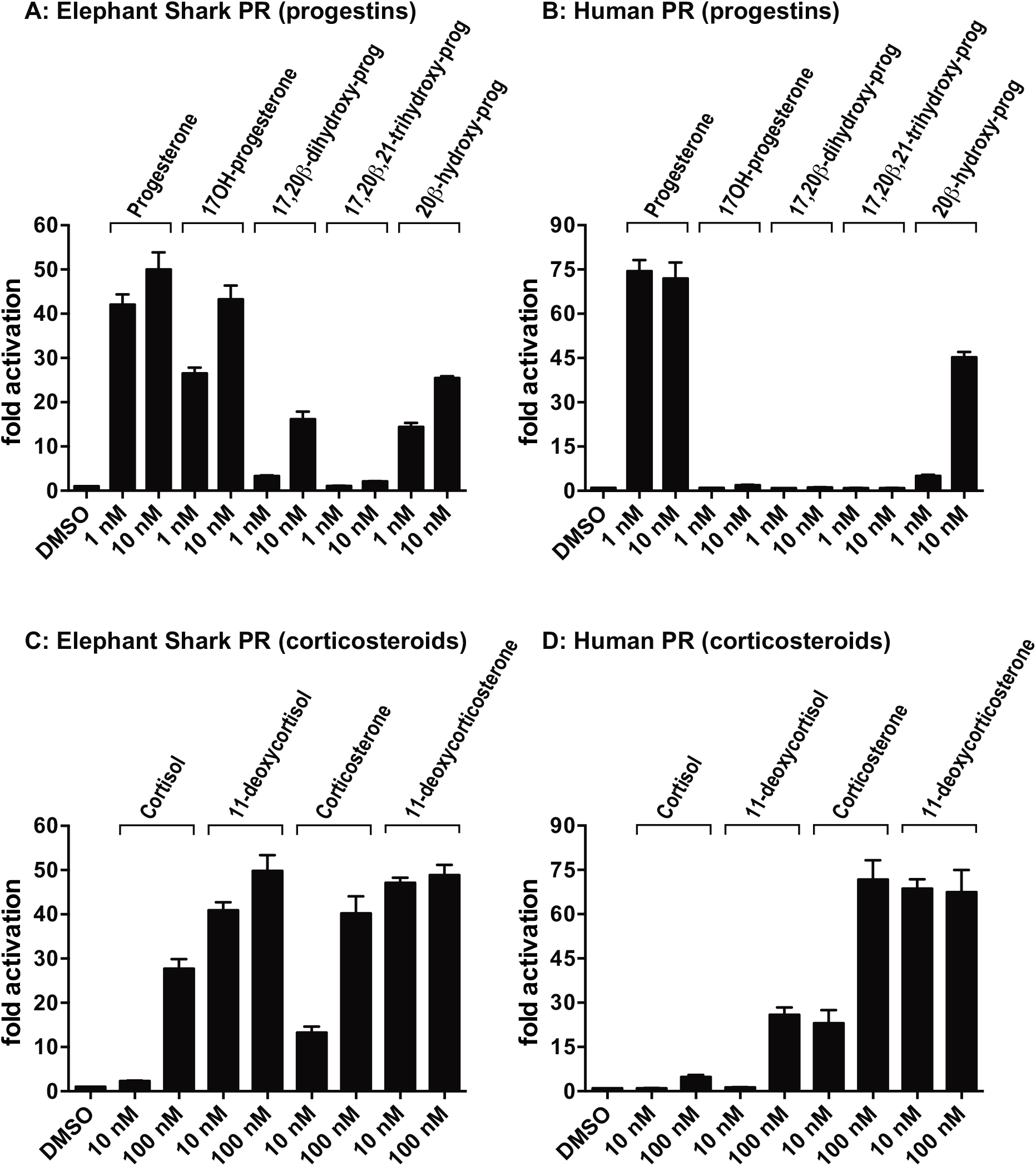
Transcriptional activation of elephant shark PR by progestins and corticosteroids. Elephant shark PR (A and C) and human PR (B and D) were expressed in HEK293 cells with an MMTV-luciferase reporter. Cells were treated with 1 and 10 nM progestins (progesterone, 17OH-progesterone, 17,20β-dihydroxy-progesterone, 17,20β,21-trihydroxy-progesterone and 20β-OH-progesterone), and 10 and 100 nM corticosteroids (cortisol, 11-deoxycortisol, corticosterone, 11-deoxycorticosterone), or vehicle alone (DMSO). Results are expressed as means ± SEM, n=3. Y-axis indicates fold-activation compared to the activity by vehicle (DMSO) alone as 1.

At 10 nM steroid, human PR responded only to progesterone and 20β-hydroxy-progesterone and not to the other progestins (Figure 3B). At 10 nM, 11-deoxycorticosterone and corticosterone activated human PR (Figure 3D).

### Concentration-dependent activation by progestins and corticosteroids of elephant shark PR and human PR

To gain a quantitative measure of progestin and corticosteroid activation of elephant shark PR and human PR, we determined the concentration dependence of transcriptional activation by progestins and corticosteroids of elephant shark PR (Figure 4A, C) and for comparison activation of human PR (Figure 4B, D). Progesterone, 17OH-progesterone, 17,20β-dihydroxy-progesterone, 20β-OH-progesterone, 11-deoxycortisol, corticosterone and 11-deoxycorticosterone activated elephant shark PR, while human PR was stimulated only by progesterone, 20β-OH-progesterone, corticosterone and 11-deoxycorticosterone, a more limited number of steroids.

**Figure 4.**
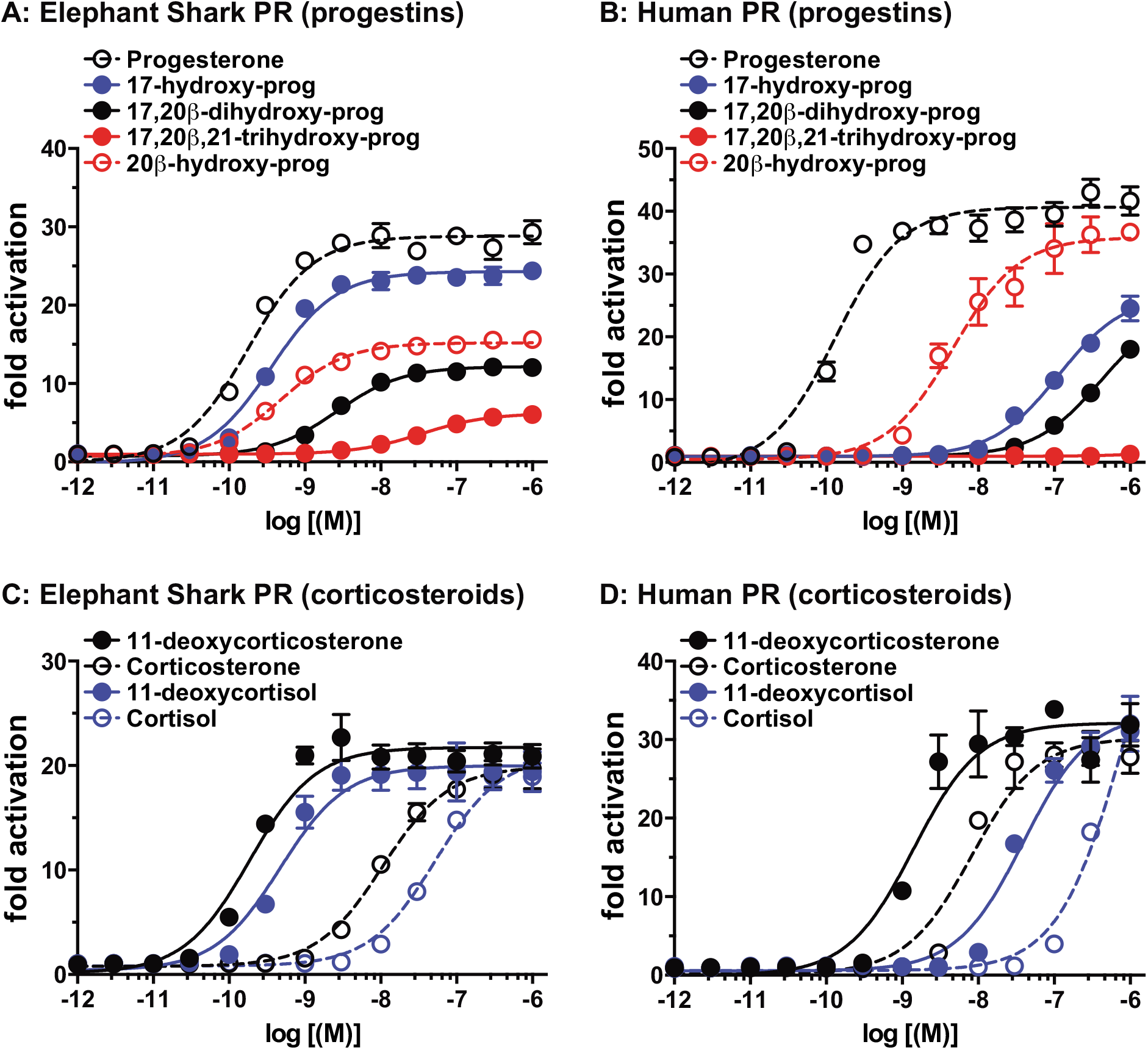
Concentration-dependent transcriptional activation of elephant shark and human PRs by corticosteroids and progestins. Elephant shark PR (A and C) and human PR (B and D) were expressed in HEK293 cells with an MMTV-luciferase reporter. Cells were treated with increasing concentrations of corticosteroids (A and B), progestins (C and D) or vehicle alone (DMSO). Y-axis indicates fold-activation compared to the activity by vehicle (DMSO) alone as 1.

Table 1 summarizes the EC50s of progestins and corticosteroids for elephant shark PR and human PR. We find that elephant shark PR has low EC50s for progesterone (0.18 nM), 17-OH-progesterone (0.36 nM), 20β-OH-progesterone (048 nM), 17α,20β-OH-progesterone (2.6 nM), 11-deoxycortisol (0.47 nM) and 11-deoxycorticosterone (0.19 nM). In contrast, human PR has low EC50s for progesterone (0.13 nM), 20β-OH-progesterone ( 4.6 nM), 11-deoxycorticosterone (1.4 nM) and corticosterone (8.2 nM).

**Table 1.** EC50 values for steroid activation of elephant shark PR and human PR.

|  | **Progesterone** | **17OH-Prog** | **20β-OH-Prog** |
| --- | --- | --- | --- |
|  | **EC50 (M)** | **EC50 (M)** | **EC50 (M)** |
| **Elephant shark PR** | **0.18 nM** | **0.36 nM** | **0.48 nM** |
| **95% confidence intervals** | **0.14-0.22 nM** | **0.29-0.45 nM** | **0.41-0.57 nM** |
| **Human PR** | **0.13 nM** | **113.0 nM** | **4.6 nM** |
| **95% confidence intervals** | **0.094-0.18 nM** | **87.3-146 nM** | **3.2-6.6 nM** |

**Fish Progestins**
|  | **17α,20β-DP** | **20β-S** |
| --- | --- | --- |
|  | **EC50 (M)** | **EC50 (M)** |
| **Elephant shark PR** | **2.6 nM** | **34.4 nM** |
| **95% confidence intervals** | **2.2-3.2 nM** | **29.6-40.2 nM** |
| **Human PR** | **408 nM** | **-** |
| **95% confidence intervals** | **342-487 nM** | **-** |

**Corticosteroids**
|  | **Cortisol** | **11-deoxycortisol** | **Corticosterone** | **DOC** |
| --- | --- | --- | --- | --- |
|  | **EC50 (M)** | **EC50 (M)** | **EC50 (M)** | **EC50 (M)** |
| **Elephant shark PR** | **52.4 nM** | **0.47 nM** | **10.5 nM** | **0.19 nM** |
| **95% confidence intervals** | **44.3-62 nM** | **0.35-0.63 nM** | **9.0-12.3 nM** | **0.13-0.26 nM** |
| **Human PR** | **826 nM** | **39.0 nM** | **8.2 nM** | **1.4 nM** |
| **95% confidence intervals** | **516-1322 nM** | **30-51 nM** | **5.5-12.1 nM** | **0.9-2.2 nM** |

Human Progestins: 17OH-Prog = 17α-hydroxy-progesterone,
20β-OH-Prog = 20β-hydroxy-progesterone,
Fish Progestins: 17α,20β-DP = 17α,20β-dihydroxy-4-pregnene-3-one,
20β-S = 17α,20β,21-trihydroxy-4-pregnen-3-one,
Corticosteroids: DOC = 11-deoxycorticosterone,

### RU486 does not inhibit transactivation of elephant shark PR

Activation of human PR by progesterone is inhibited by RU486 (29,30,33). Indeed, at 0.1 nM and 1 nM, RU486 inhibits of activation by 1 nM progesterone of human PR (Figure 5A). Benhamou et al. (35) reported that Gly-722 in human PR, is essential for the inhibition of progesterone activation of human PR by of RU486. We confirm that RU486 does not inhibit the human PR Cys722 mutant (Figure 5C).

**Figure 5.**
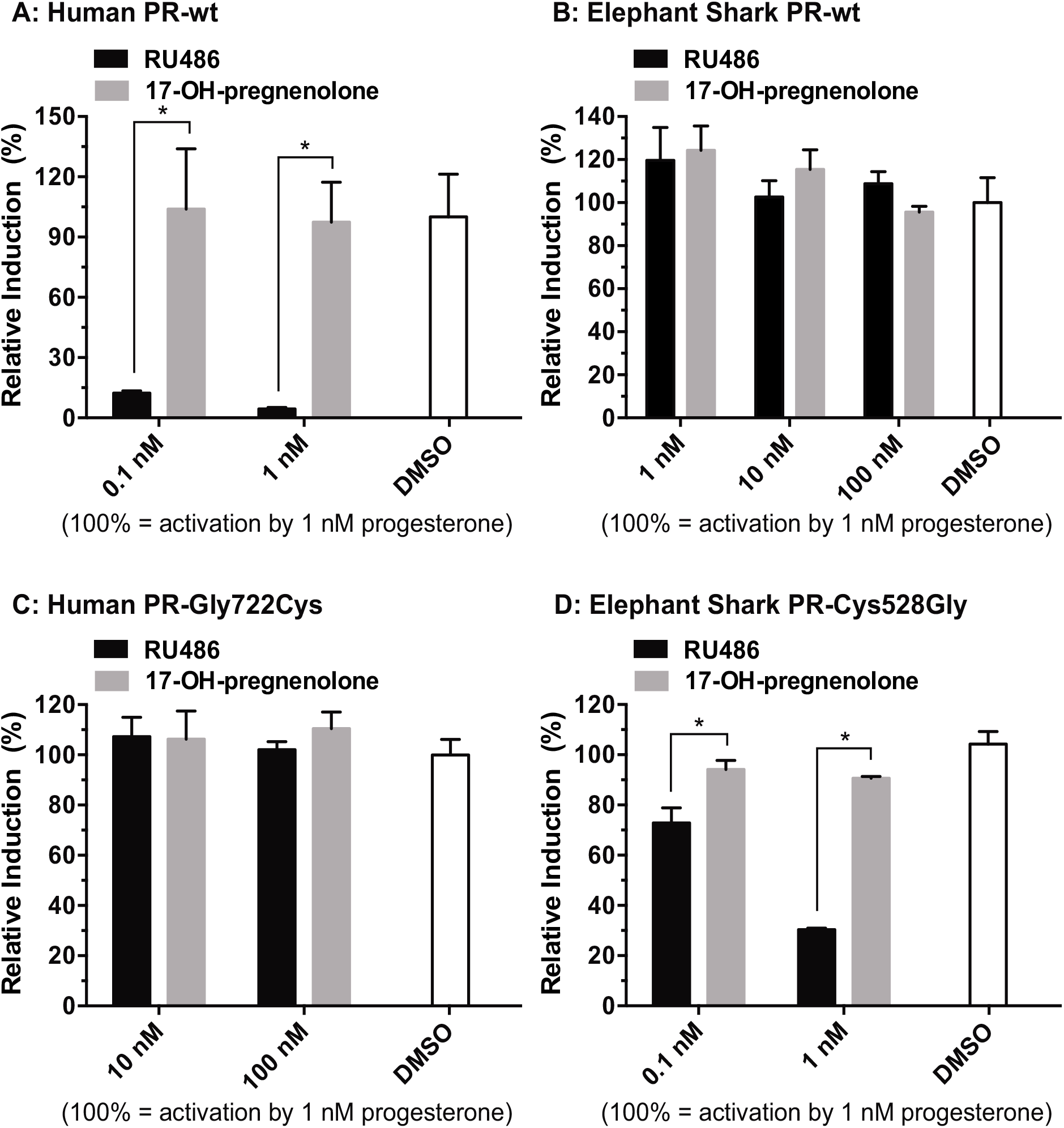
Effect of RU486 for Prog-induced activation of PR. Wild-type of human PR (A) or elephant shark PR (B) was expressed in HEK293 cells with an MMTV-luciferase reporter. Cells with human PR were treated with 1 nM progesterone and either 0.1 nM or 1 nM RU486, 17OH-pregnenolone or DMSO. Cells with elephant shark PR were treated with 1 nM progesterone and with either 1 nM, 10 nM or 100 nM RU486, 17OH-pregnenolone or DMSO. Human PR-Gly722Cys (C) or elephant shark PR-Cys528Gly (D) was expressed in HEK293 cells with an MMTV-luciferase reporter. Cells with human PR-Gly722Cys were treated with 1 nM progesterone and with either 10 nM or 100 nM RU486, 17OH-pregnenolone or DMSO. Cells with elephant shark PR-Cys528Gly were treated with 1 nM progesterone and with either 0.1 nM or 1 nM RU486 or 17OH-Pregnenolone Relative inductions were normalized between 0 and 100%, where 0 and 100 were defined as the bottom and tip value in vehicle-treated and 1 nM progesterone treated, respectively. Results are expressed as means ± SEM, n=3. * *P* < 0.05 compared with vehicle treatment (student’s *t*-test).

Cys-528 of elephant shark PR corresponds to Gly-722 of human PR, which predicts that RU486 would not inhibit progesterone activation of wild type elephant shark PR, and indeed, as shown in Figure 5B, activation by 1 nM progesterone of elephant shark PR was not inhibited by 100 nM RU486. As expected, RU486 inhibits progesterone activation of elephant shark PR Gly528 (Figure 5D).

### Steroid activation of human PR Gly722Cys and elephant shark PR Cys528Gly

To search for a biological basis for the functional changes in human PR due to Gly-722 in human PR we constructed a human PR-Cys722 mutant and an elephant shark PR-Gly528 mutant and studied their activation by various progestins and corticosteroids (Figure 6). Elephant shark PR-Gly528 had a weaker response to 11-deoxycortisol and 17-hydroxy-progesterone. Human PR-Cys722 displayed increased activation by 11-deoxycortisol and decreased activation by corticosterone.

**Figure 6.**
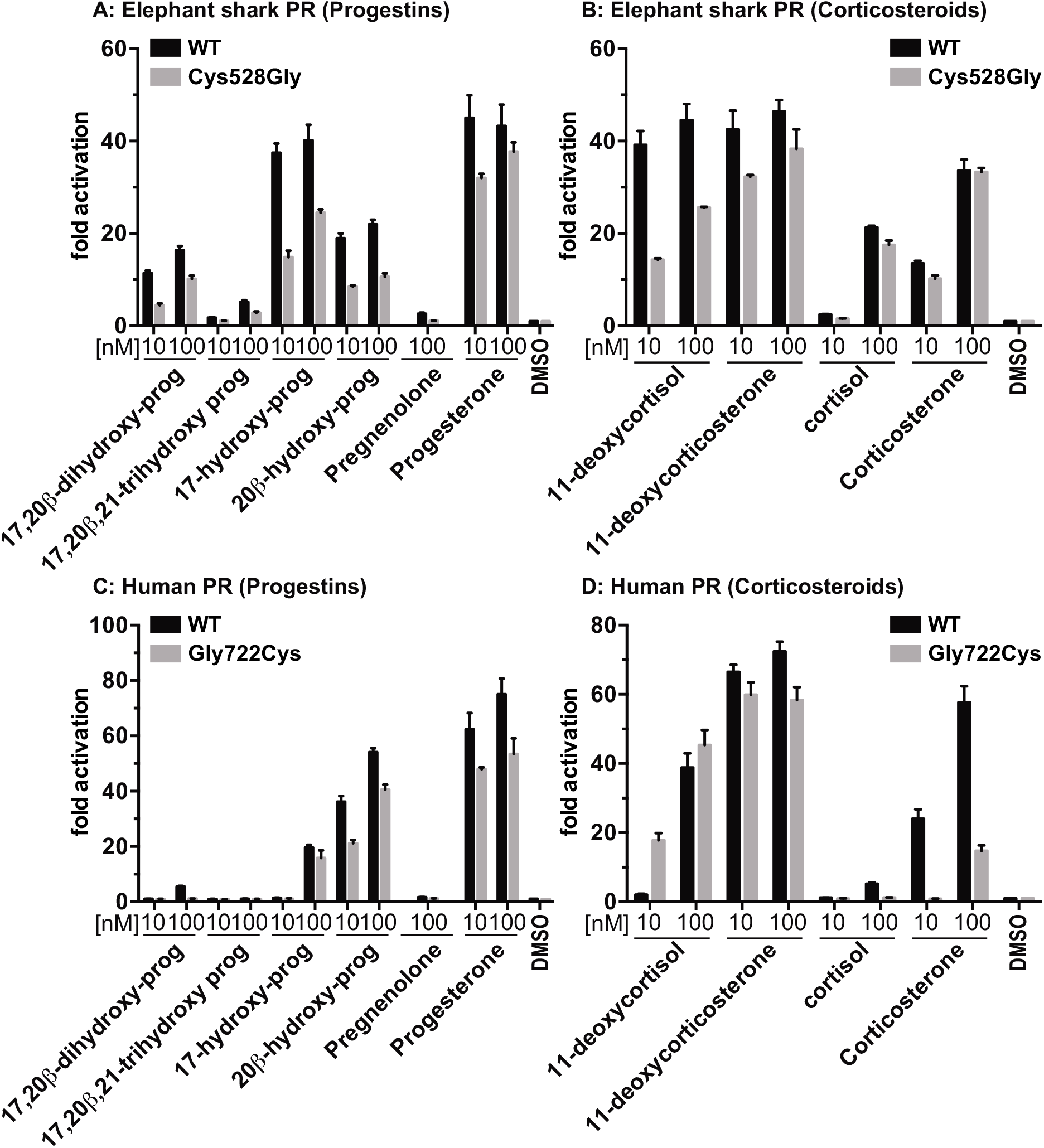
Progestin and corticosteroid activation of elephant shark PR-Cys528Gly and human PR-Gly722Cys. Elephant shark PR (A and C), and human PR (B and D) were expressed in HEK293 cells with an MMTV-luciferase reporter. Cells transfected with PRs were treated with increasing concentrations of Prog or vehicle alone (DMSO) (A and B). Cells were treated with 10 nM progestins (Prog, 17OH-Progesterone, pregnenolone 17,20β-dihydroxy-progesterone, 17,20β,21-trihydroxy-progesterone, 20β-OH-progesterone), corticosteroids (cortisol, 11-deoxycortisol, corticosterone, 11-deoxycorticosterone), or vehicle alone (DMSO) (C and D). Results are expressed as means ± SEM, n=4. Y-axis indicates fold-activation compared to the activity of control vector with vehicle (DMSO) alone as 1.

### Concentration-dependent transcriptional activation of elephant shark PR-Cys528Gly and human PR-Gly722Cys

To gain a quantitative measure of progestin and corticosteroid activation of the cysteine-528 to glycine-528 mutation in elephant shark PR and the glycine-722 to cysteine-722 mutation in human PR, we determined the concentration dependence of transcriptional activation by progestins and corticosteroids of Cys528Gly elephant shark PR (Figure 7A, C) and for comparison activation of Gly722Cys human PR (Figure 7B, D). In a separate experiment we investigated the response of Cys528Gly elephant shark PR and Gly722Cys human PR to corticosterone (Figure E, F).

**Figure 7.**
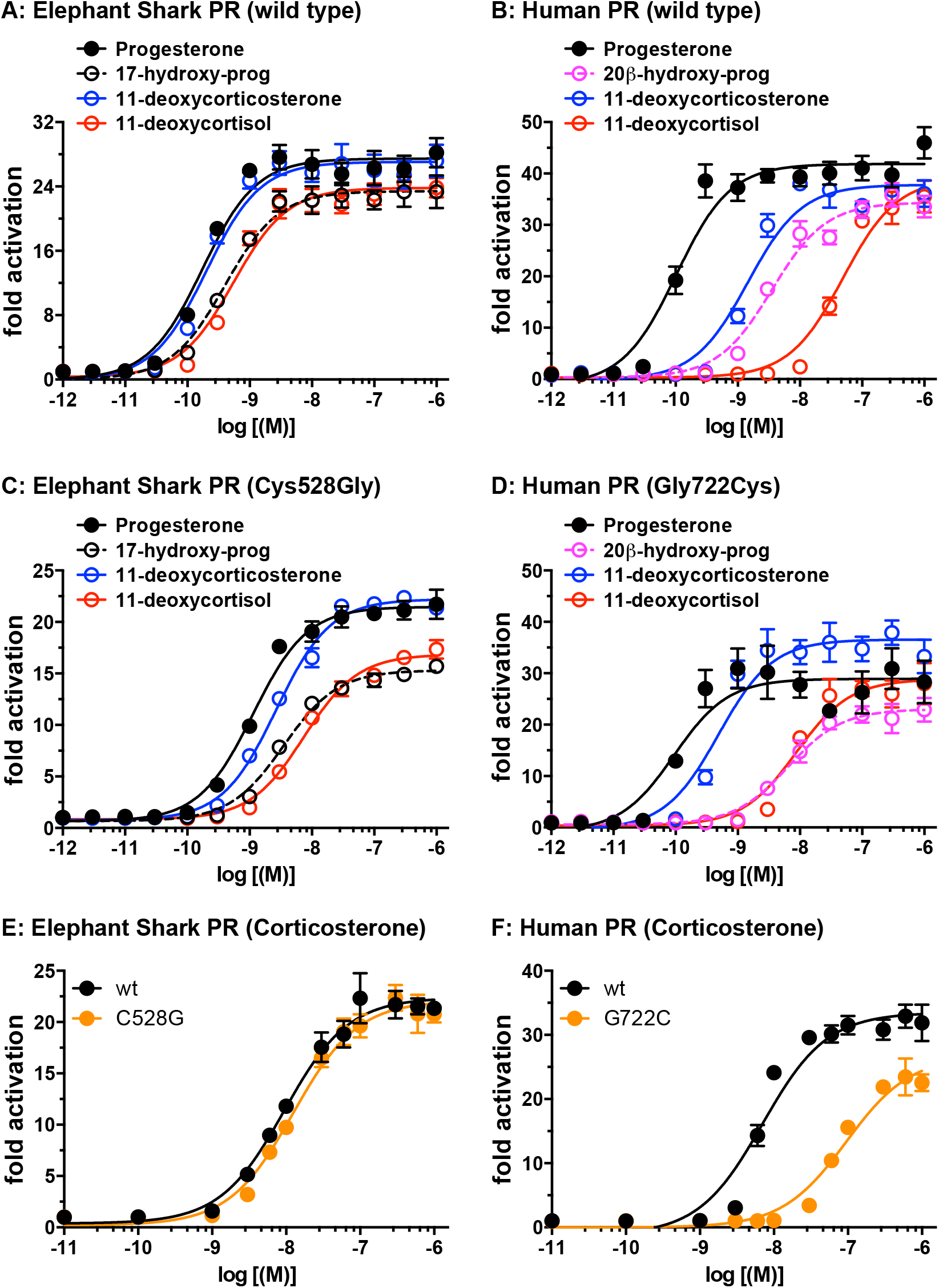
Concentration dependent transcriptional activation by progestins and corticosteroids of elephant shark PR-Cys528Gly and human PR-Gly722Cys. Elephant shark PR (A, C and E), and human PR (B, D and F) were expressed in HEK293 cells with an MMTV-luciferase reporter. Cells transfected with PRs were treated with increasing concentrations of progesterone, 20β-OH-progesterone, 11-deoxycortisol, 11-deoxycorticosterone or corticosterone. Results are expressed as means ± SEM, n=4. Y-axis indicates fold-activation compared to the activity of control vector with vehicle (DMSO) alone as 1.

Table 2 shows the EC50s calculated from the curves in Figure 7. Progesterone, 20β-OH-progesterone, 11-deoxycortisol, corticosterone and 11-deoxycorticosterone activated elephant shark PR, while human PR was stimulated only by progesterone, 20β-OH-progesterone, corticosterone and 11-deoxycorticosterone, a more limited number of steroids. One or more of these changes may have been selective for the evolution an ancestor of glycine-722 in a PR in an ancestral platypus at the base of the mammalian line.

**Table 2.** EC50 values for steroid activation of wild-type and Cys528Gly mutant elephant shark PR and Gly722Cys mutant human PR.

**Progestins**
|  | **Progesterone** | **17OH-Prog** | **20β-OH-Prog** |
| --- | --- | --- | --- |
|  | **EC50 (M)** | **EC50 (M)** | **EC50 (M)** |
| **Elephant shark PR** | **0.17 nM** | **0.4 nM** | **-** |
| **95% confidence intervals** | **0.13-0.24 nM** | **0.29-0.56 nM** | **-** |
| **Elephant shark PR-Cys528Gly** | **1.1 nM** | **3.4 nM** | **-** |
| **95% confidence intervals** | **0.92-1.4 nM** | **2.8-4.0 nM** | **-** |
| **Human PR** | **0.1 nM** | **-** | **3.5 nM** |
| **95% confidence intervals** | **0.07-0.15 nM** | **-** | **2.6-4.7 nM** |
| **Human PR-Gly722Cys** | **0.095 nM** | **-** | **6.4 nM** |
| **95% confidence intervals** | **0.05-0.2 nM** | **-** | **4.4-9.3 nM** |

**Corticosteroids**
|  | **DOC** | **11-deoxycortisol** | **Corticosterone** |
| --- | --- | --- | --- |
|  | **EC50 (M)** | **EC50 (M)** | **EC50 (M)** |
| **Elephant shark PR** | **0.2 nM** | **0.55 nM** | **9.4 nM** |
| **95% confidence intervals** | **0.14-0.28 nM** | **0.41-0.74 nM** | **7.0-12.7 nM** |
| **Elephant shark PR-Cys528Gly** | **2.7 nM** | **7.4 nM** | **12.5 nM** |
| **95% confidence intervals** | **2.3-3.1 nM** | **6.2-8.9 nM** | **9.4-16.5 nM** |
| **Human PR** | **1.4 nM** | **49.2 nM** | **7.0 nM** |
| **95% confidence intervals** | **1.0-2.1 nM** | **36-68 nM** | **4.8-10.2 nM** |
| **Human PR-Gly722Cys** | **0.5 nM** | **8.7 nM** | **95.0 nM** |
| **95% confidence intervals** | **0.32-0.76 nM** | **5.7-13.4 nM** | **67-134 nM** |

**Progestins: 17OH-Prog = 17α-hydroxy-progesterone, 20β-OH-Prog = 20β-hydroxy-progesterone.**

## Discussion

An ortholog of human PR along with the corticoid receptor (CR), the ancestor of the MR and GR, first appears in the more ancient cyclostomes (jawless fish), which has descendants in modern lamprey and hagfish (3,27,42). It is in cartilaginous fish that distinct orthologs of human MR and GR first appear (43–45) along with the beginning of the evolution of differences in the responses of the PR, MR and GR for progestins and corticosteroids that appear in terrestrial vertebrates and ray-finned fish (16,17,24,40,41,43–52).

Here we report that elephant shark PR has a strong response to progesterone (EC50 0.18 nM) and 17-OH-progesterone (EC50 0.36 nM), as well as to 11-deoxycorticosterone (EC50 0.19 nM) (Figure 4, Table 1), a corticosteroid with close structural similarity to progesterone (Figure 1). Elephant shark PR also is activated by 20β-OH-progesterone (EC50 0.48 nM) and 17,20β-dihydroxy-progesterone (EC50 2.6 nM) and 11-deoxycortisol (EC50 0.47 nM). This broad response to steroids contrasts with the selectivity of human PR, which has a strong response to progesterone (EC50 0.13 nM) and 11-deoxycorticosterone, (EC50 1.4 nM) and a weaker response to corticosterone, (EC50 8.2 nM). The advantage this selectivity of human PR is not known.

The evolution of the response of human PR to RU486 is intriguing because RU486 is not a physiological ligand for the PR. The glycine-722 in human PR (Figure 2) that confers antagonist activity for RU486 towards human PR first appears in platypus, a basal mammal (Figure 2). Human PR with Cys722, to mimic an ancestral PR, had increased activation by 11-deoxycortisol and decreased activation by corticosterone.

**Figure 2.**
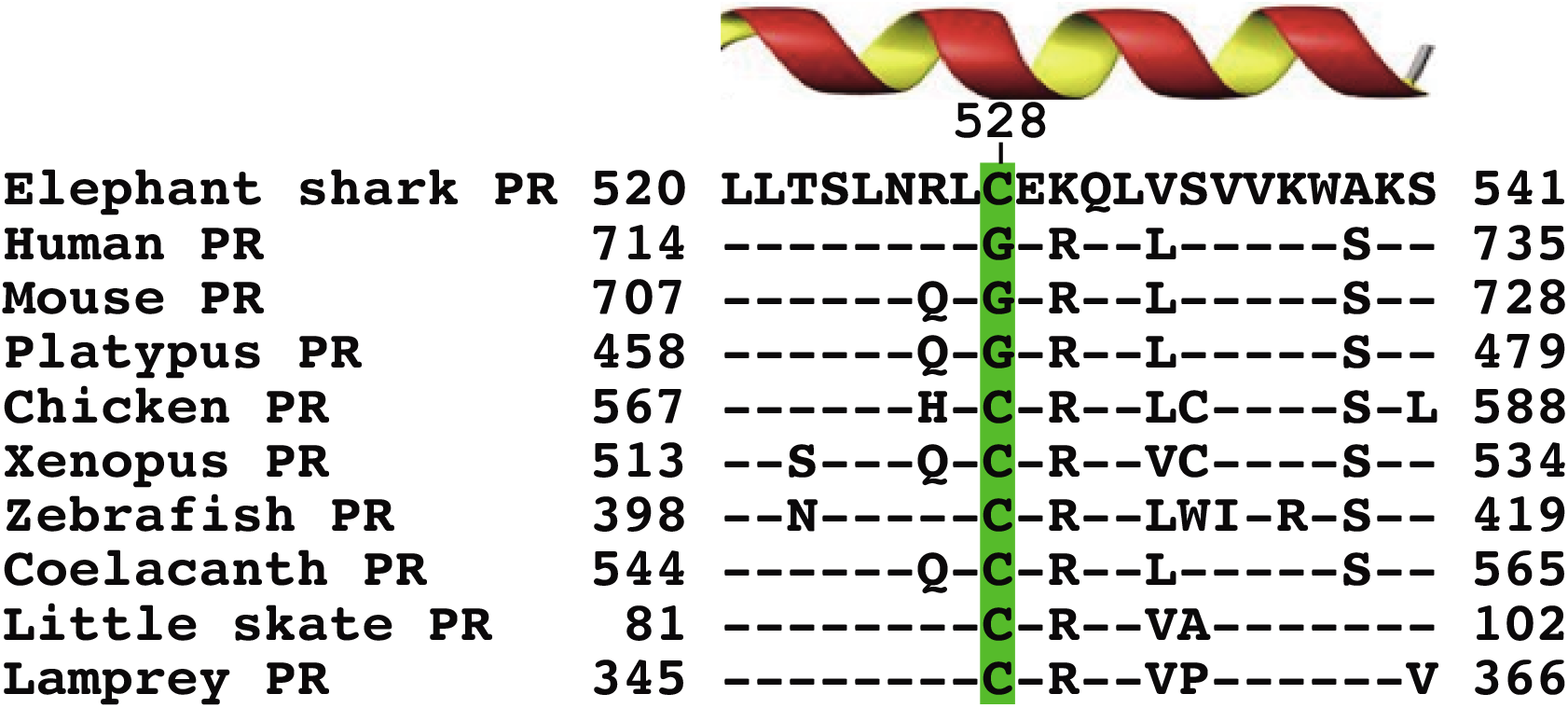
Alignment of α helix-3, containing a key amino acid necessary for RU486 inhibition of human PR and activation of elephant shark PR. Alignment of α-helix-3 in human PR, containing Gly-722 that is essential for RU486 inhibition of progesterone activation of human PR (35), the PR in elephant shark and other selected vertebrates. RU486 activates elephant shark PR, which contains cysteine-528 corresponding to human PR Gly-722. A glycine first appears in this position in platypus PR, a basal mammal. Amino acids that are identical to amino acids in elephant shark PR are denoted by (−).

The lower activation of elephant shark PR by 17,20β-dihydroxy-progesterone compared to progesterone is intriguing because in zebrafish (16,17), this is reversed with higher activity for 17,20β-dihydroxy-progesterone compared to progesterone indicating that during the evolution of ray-finned fish the response of the PR to progesterone diminished and the response to 17,20β-dihydroxy-progesterone increased (21,22,36,37). The absence of progesterone as a ligand for ray-finned fish PR may be relevant for progesterone functioning as a ligand for fish MR (39,41,45,51,53).

## Funding, Contributions and Competing Interests

### Funding

This work was supported by Grants-in-Aid for Scientific Research from the Ministry of Education, Culture, Sports, Science and Technology of Japan (19K067309 to Y.K.), and the Takeda Science Foundation (to Y.K.). M.E.B. was supported by Research fund #3096.

### Author contributions

X.L. and Y.K. carried out the research and analyzed data. W.T. and S.H. aided in the collection of animals. S.I. provided steroids used in this study. Y.K. and M.E.B. conceived and designed the experiments. X.L., Y.K. and M.E.B. wrote the paper. All authors gave final approval for publication.

### Competing Interests

We have no competing interests.

## Notes

### Competing Interest Statement

The authors have declared no competing interest.

### Summary of Updates

Analysis of concentration dependent of steroid activation of Cys528Gly mutant elephant shark PR and Gly722Cys mutant human PR has been added.

